# ProphET, Prophage Estimation Tool: a standalone prophage sequence prediction tool with self-updating reference database

**DOI:** 10.1101/176750

**Authors:** João L. Reis-Cunha, Daniella C. Bartholomeu, Ashlee M. Earl, Bruce W. Birren, Gustavo C. Cerqueira

## Abstract

Prophages are a significant force in prokaryote evolution. The remaining sequences of a bacteriophage integration event are known for altering gene expression, enabling creative destruction of the bacterial genome and to induce pathogenicity by harboring and transposing virulence and antibiotic resistance factors. In the light of the dreadful expansion of antibiotic resistance bacteriophages have gathered renewed interest from the scientific community and public health decision makers as a promising long forgotten alternative to control bacterial infections. Cataloging the repertoire of prophages and their integration sites is an important initial step in the understanding of bacteriophages either as tool or as a threat. In this work, we present ProphET (Prophage Estimation Tool), a standalone application without the limitations of their web based counterparts and which identifies prophages in bacterial genomes with higher precision than similar applications.

## Introduction

Prophages are a significant force in prokaryote evolution. The remaining sequences of a bacteriophage integration event are known for altering gene expression, enabling creative destruction of the bacterial genome and to induce pathogenicity by harboring and transposing virulence and antibiotic resistance factors (Aziz *et al.*, 2005; Casjens, 2003; Laing *et al.*, 2012; Yan *et al.*, 2015). In the light of the dreadful expansion of antibiotic resistance bacteriophages have gathered renewed interest from the scientific community and public health decision makers as a promising long forgotten alternative to control bacterial infections. Cataloging the repertoire of prophages and their integration sites is an important initial step in the understanding of bacteriophages either as tool or as a threat and with this aim a handful of prophage identification tools have been designed (Hendrix, 2002; Suttle, 2007; Aziz *et al.*, 2005). However, the prediction of prophages in bacterial genomes is hampered by their low similarity across families and very distinct genome sizes (Pope *et al.*, 2015; Grose and Casjens, 2014). Among the few publicly available applications (Fouts, 2006; Akhter *et al.*, 2012) Prophinder and PHAST are the most well-known (Zhou *et al.*, 2011; Lima-Mendez *et al.*, 2008). Both programs are only available as web applications, which limits their throughput and routinely burdens users with long response times during periods of high demand. This is less than ideal for users affiliated with large genomic institutes, where generally a single project consists of the sequencing and the analysis of hundreds of bacterial genomes. In those cases, a prophage prediction tool that can be executed locally and without the constraints associated with web based tools is the most appropriate solution. In this work, we present ProphET (Prophage Estimation Tool), a standalone application without the limitations of their web based counterparts and which identifies prophages in bacterial genomes with higher precision than PHAST and Prophinder. ProphET application is available from: github.com/jaumlrc/ProphET

## Application description

### Self-updating Database

ProphET predictions relies on similarity searches against a database of prophage genes created automatically during Prophet’s first execution and by default updated monthly. Database self-updates can be either turned off or forced via command parameters.

ProphET database consists of the proteome sequences of all available phage genomes (NCBI Genbank) belonging to 18 families listed in Krupovic 2011 (Krupovic *et al.*, 2011). The current version contains 142,575 protein sequences representing the proteome of 1,435 bacterial and archaeal, DNA and RNA bacteriophages, with linear, circular or segmented genomes. However, as novel bacteriophage genomic sequences are deposited in Genbank the self-update module will add those to ProphET’s database and those numbers will increase. Protein sequences of ABC transporters were excluded from the database, as they are highly conserved in prokaryotes and eukaryotes, and could lead to false-positive phage predictions (Davidson *et al.*, 2008).

### Input Files

ProphET takes as input two files: the bacterial genomic sequence in FASTA format and its correspondent gene annotation as a GFF file.

### Phage prediction

ProphET identifies prophage based on the nucleotide density of phage-like genes, a process that can be divided in three steps: similarity search, prophage density and polishing.

### Similarity search

The coding sequence of each gene is extracted from the bacterial genomic sequence according to the coordinates in the user provided GFF file and then translated into protein sequences. Phage-like genes are identified by a BLASTP search of the bacterial conceptual proteome against ProphET reference database. Only matches with an e-value lower than 10^-5^ are considered.

### Prophage density

A sliding window of length 10 kb runs through the bacterial genome in increments of 1 kb. Instances with at least 8 phage-like genes with an overall length longer than half of the length of the sliding window (5 Kb) are selected (Supplemental Figure S1). Overlapping selected windows are merged in larger segments and assigned as preliminary prophage predictions

The length of the sliding window and the minimum number of phage-like genes in each window were defined according to distribution of those traits for the bacteriophage genomes in ProphET database: only 6.2% of genomes are shorter than 10kb and those have less than 16 genes.

### Polishing

Next, ProphET trims the borders of preliminary prophage prediction at the last phage-like gene. As tRNA genes are hotspots of prophage integration (Campbell, 2003), our program searches the 3 kb upstream and downstream of the putative prophage borders for tRNA genes. If a tRNA gene is found, it extends or contracts the prophage predicted border accordingly, resulting in the final prophage prediction (Supplemental Figure S2).

## Performance and comparison with other predictors

The performance of ProphET was evaluated and compared to other prophage prediction tools using a dataset of 54 bacterial genomes containing 287 prophages manually annotated by Casjens (Casjens, 2003). The same dataset was used as the gold standard in the articles reporting the performance of Prophinder v0.4 (Lima-Mendez *et al.*, 2008) and PHAST (Zhou *et al.*, 2011). We considered as true positives (TP) the overall length of the overlap between prophage predictions and the gold standard; false positives (FP) as the overall length of predictions that do not overlap with the gold standard; false negatives (FN) as the overall length of regions of the gold standard that were not included in the predictions (Supplemental Figure S3). The sensitivity (TP/TP+FN) and positive predictive value or PPV (TP/TP+FP) were computed and compared to those obtained by Prophinder and PHAST.

Based on this analysis ProphET had a higher overall performance among the three programs, with a sensitivity of 0.76 and a PPV of 0.85, while PHAST had 0.79 sensitivity and 0.72 PPV and Prophinder had 0.71 sensitivity and 0.75 PPV, under strict settings. The slightly higher sensitivity obtained by PHAST (delta=0.03) when compared with ProphET comes with the cost of a great loss in PPV (delta=0.13). The higher sensitivity obtained by PHAST may not be an accurate representation of its performance in regular conditions, as PHAST reference database includes the gold standard (Casjens, 2003) (Srividhya 2006).

Some predictions were corroborated by more than one application but were not validated by the gold standard. Those may either represent real phages that were unknown when the gold standard was published or a common flaw shared by more than one predictor that needs to be further addressed. The majority of the prophages missed by ProphET were neither detected by the other predictors. Those consist of short prophages without phage-like genes, according to our reference database. This limitation may be addressed as more phages are annotated and automatically included in our reference database.

**Figure 1.**
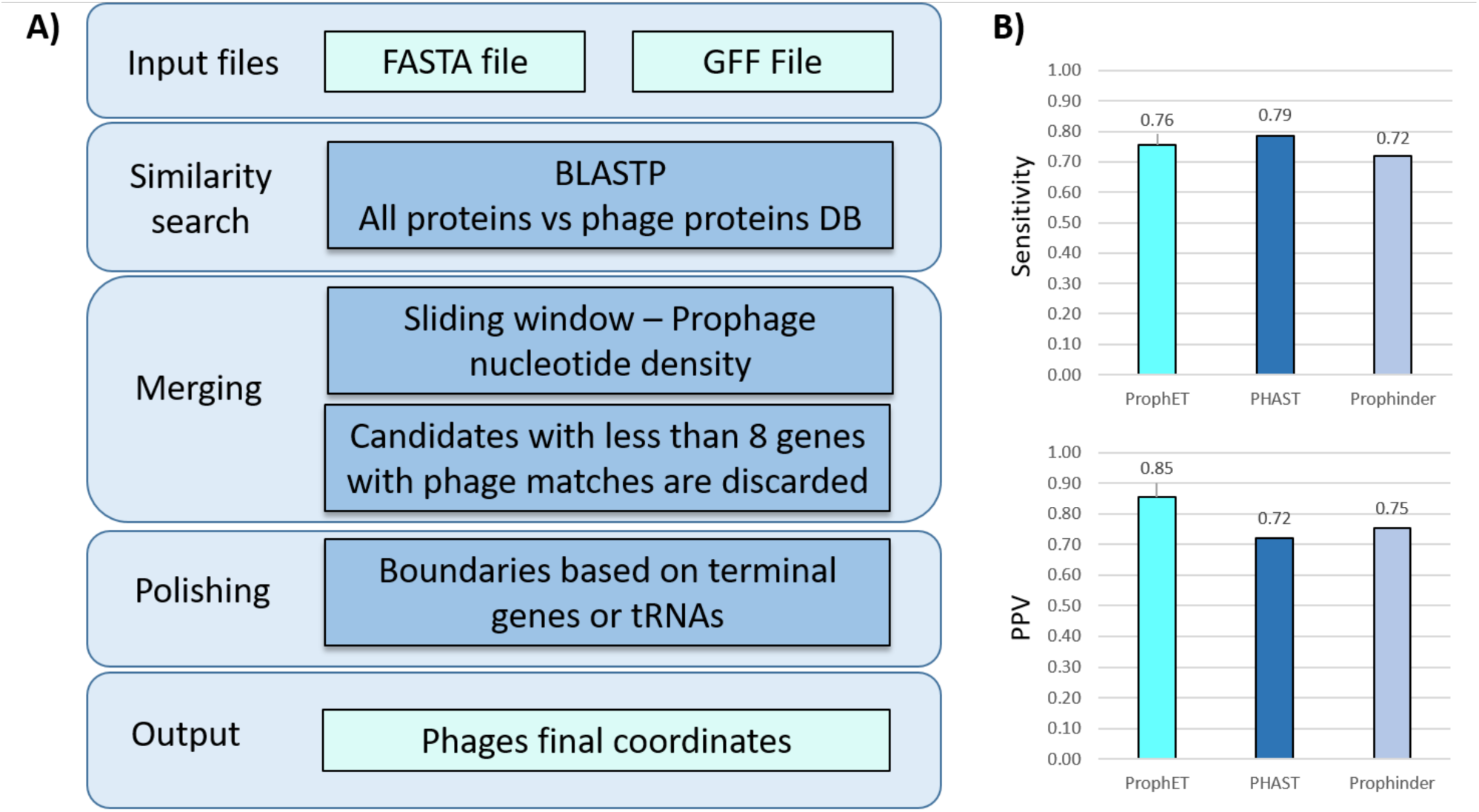
ProphET pipeline and prophage prediction performance: **(A)** ProphET requires as input the bacterial genomic sequence (FASTA file) and the corresponding coordinates of protein coding genes and tRNA (GFF files). The conceptual proteome of bacterial genome is automatically generated and submitted to a BLASTP search against ProphET phage database. Significant matches are assigned as phage-like genes. A sliding window approach is then used to identify regions with high-density of phage-like genes. Finally, the coordinates of predicted prophage are defined according to the boundaries of the last phage-like gene or tRNA genes flanking the high-density phage-like genes region. **(B)** Comparison of the performance of Prophinder (light-blue), PHAST (dark-blue) and ProphET (cyan) assessed in terms sensitivity and PPV.

## Supplemental material

**Supplemental figure S1.**
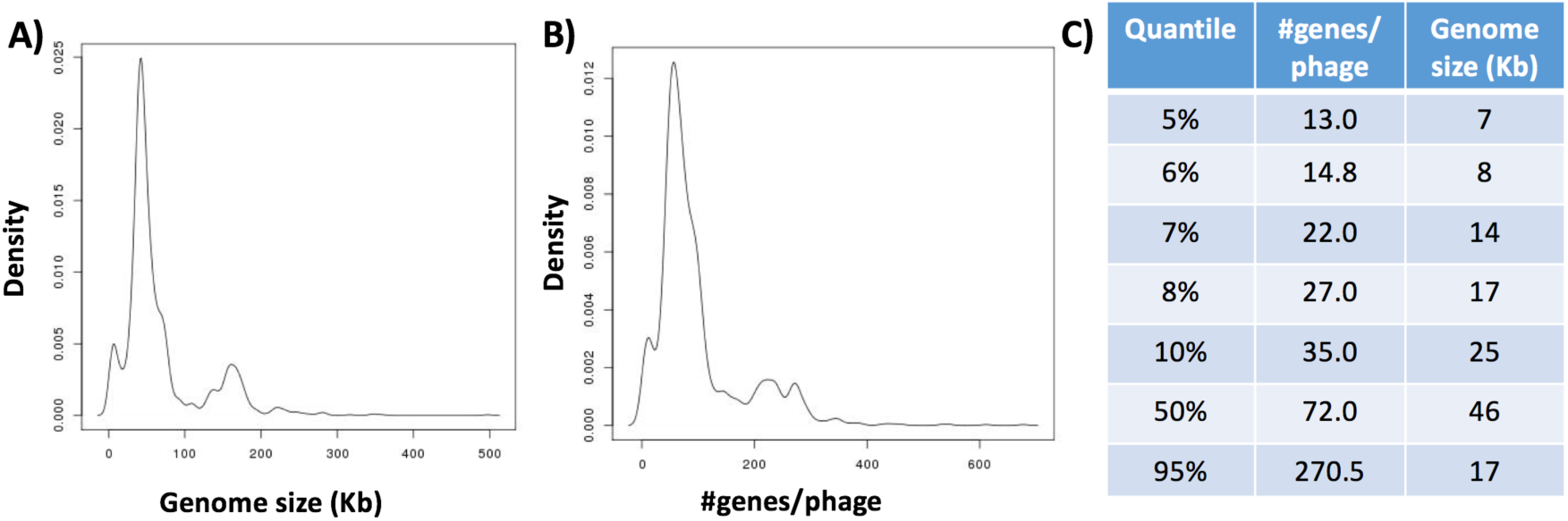
Distribution of phage sizes and the number of genes per phage in ProphET reference database. Histogram of phage genome size **(A),** gene density **(B)** and the most relevant quartiles **(C)** among the 1,535 phages in ProphET database.

**Supplemental figure S2.**
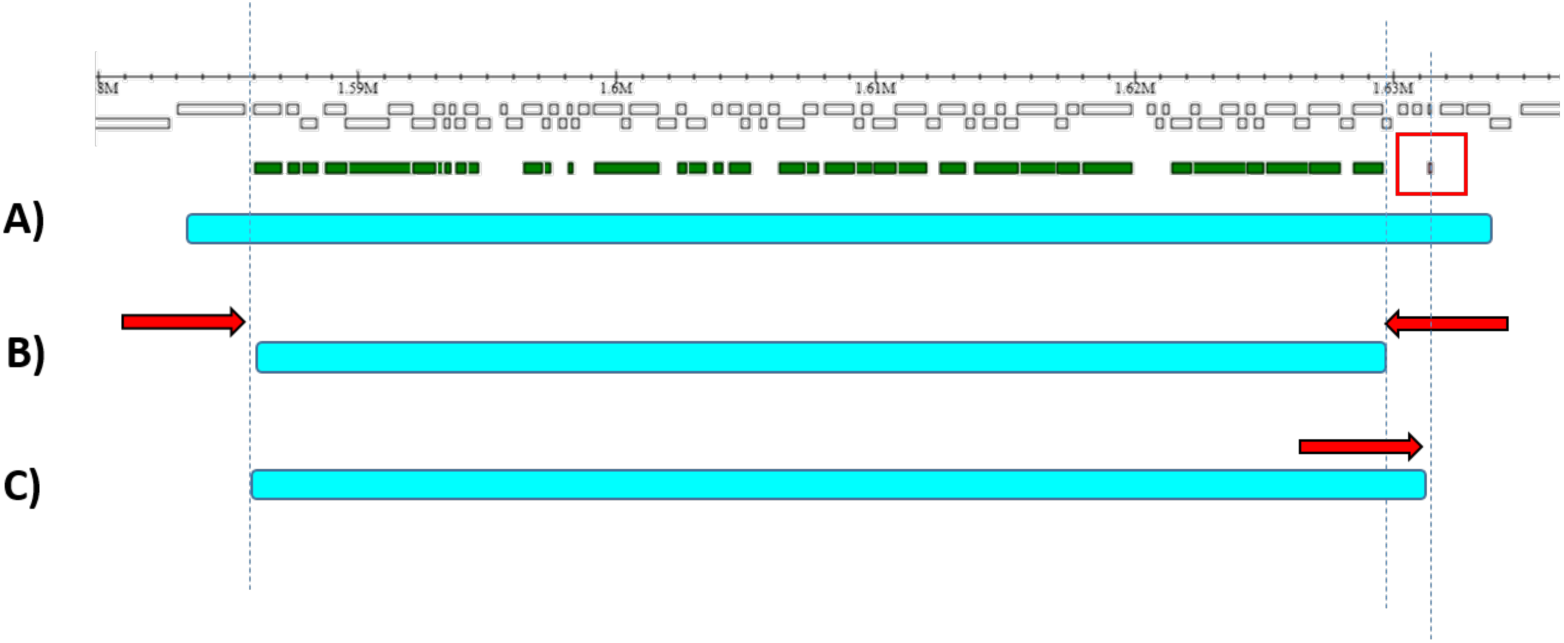
ProphET polishing stage. Ruler on top indicates the coordinates of preliminary prediction of a prophage in a bacterial genome, white boxes correspond to annotated genes and green boxes to phage-like genes as assigned by ProphET. Cyan horizontal bars depict the iterative adjustments of the boundaries of the predicted prophage. After the preliminary prophage predictions is defined **(A)**, borders are trimmed to the first and last phage-like gene within those boundaries **(B)**. Next, ProphET searches for tRNA genes within a 3 kb region upstream and downstream of the predicted prophage ends. If those are found, phage borders are extended or contracted to match the outwards coordinates of the tRNA genes **(C)**.

**Supplemental figure S3.**
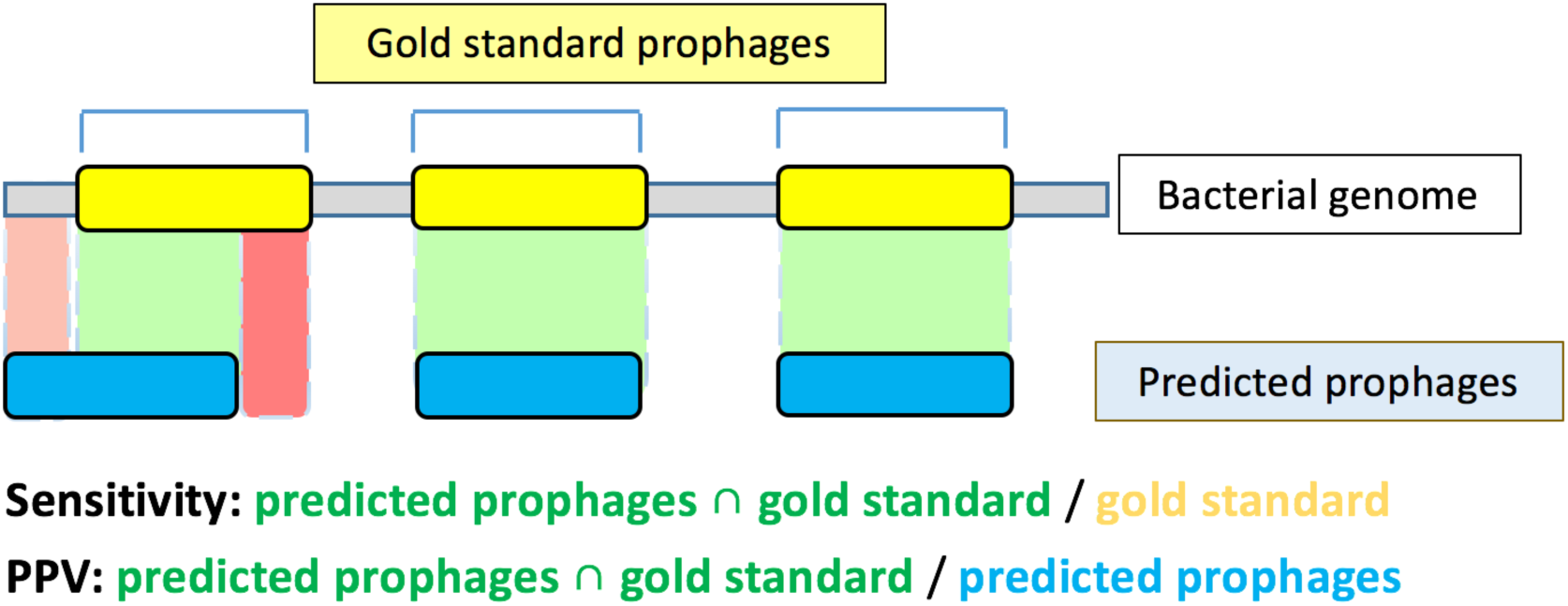
Sensitivity and PPV. Gray line corresponds to a bacterial genome, the yellow and blue boxes corresponds respectively to prophages in the gold standard and in the prediction. The sensitivity consists of the length of the intersection between prophages in the gold standard and predicted prophages (regions in green) divided by the length of prophages in the gold standard. PPV consist of the same intersection divided by the length of predicted prophages. Light and dark red areas indicate respectively false positives and false negatives.

